# Cotranscriptional demethylation induces global loss of H3K4me2 from active genes in *Arabidopsis*

**DOI:** 10.1101/2023.02.17.528985

**Authors:** Shusei Mori, Satoyo Oya, Mayumi Takahashi, Kazuya Takashima, Soichi Inagaki, Tetsuji Kakutani

## Abstract

Based on studies of animals and yeasts, methylation of histone H3 lysine 4 (H3K4me1/2/3, for mono-, di-, and tri-methylation, respectively) is regarded as the key epigenetic modification of transcriptionally active genes. In plants, however, H3K4me2 correlates negatively with transcription, and the regulatory mechanisms of this counterintuitive H3K4me2 distribution in plants remain largely unexplored. A previous genetic screen for factors regulating plant regeneration identified Arabidopsis LYSINE-SPECIFIC DEMETHYLASE 1-LIKE 3 (LDL3), which is a major H3K4me2 demethylase. Here, we show that LDL3-mediated H3K4me2 demethylation depends on the transcription elongation factor Paf1C and phosphorylation of the C-terminal domain (CTD) of RNA polymerase II (RNAPII). In addition, LDL3 binds to phosphorylated RNAPII. These results suggest that LDL3 is recruited to transcribed genes by binding to elongating RNAPII and demethylates H3K4me2 cotranscriptionally. Importantly, the negative correlation between H3K4me2 and transcription is disrupted in the *ldl3* mutant, demonstrating the genome-wide impacts of the transcription-driven LDL3 pathway to control H3K4me2 in plants. Our findings implicate H3K4me2 in plants as chromatin memory for transcriptionally repressive states, which ensures robust gene control.

## Introduction

In eukaryotes, transcription and histone modifications influence each other, generating a basis of epigenetic memory (Krogan et al. 2003; Ng et al. 2003; Soares et al. 2017; Holoch et al. 2021; Inagaki et al. 2010). Methylation of histone H3 lysine 4 (H3K4me) is one of the highly conserved histone modifications found in actively transcribed genes. H3K4me occurs in three (different) forms: mono-, di-, and tri-methylation (H3K4me1/2/3, respectively). H3K4me3 and H3K4me2 accumulate predominantly in regions near the transcription start site (TSS), whereas H3K4me1 is distributed within transcribed regions. This pattern is conserved among eukaryotes, including yeast (Pokholok et al. 2005), animals (Barski et al. 2007), and plants (Zhang et al. 2009; Oya et al. 2022). H3K4me is widely considered an active mark even though this concept is still under debate (Howe et al. 2017; Henikoff and Shilatifard 2011; Oya et al. 2022). Indeed, while H3K4me3 is strongly correlated with the transcription level in many species of fungi, animals, and plants (Barski et al. 2007; Howe et al. 2017; Oya et al. 2022; Zhang et al. 2009), correlations with transcription levels vary for H3K4me2 and H3K4me1. Intriguingly, H3K4me2 is negatively correlated with transcription levels in plants (Liu et al. 2019). This counterintuitive observation is consistent with other studies that suggest that H3K4me2 acts as a repressive mark (Oya et al. 2022; Liu et al. 2019; Wang et al. 2022) An important question is how the negative correlation between H3K4me2 and transcription is established, presumably through bidirectional interactions between transcription machinery and regulators of H3K4me2.

The distributions of H3K4me1/2/3 should be determined by H3K4 methyltransferases, as well as demethylases, most likely through interaction with other chromatin components, including the transcription machinery. An extensively studied example is the interaction between the transcription machinery and the yeast H3K4 methyltransferase Set1. Set1 binds transcription machinery (Ng et al. 2003; Bae et al. 2020), and recruitment of the methyltransferase to RNA polymerase II is required for histone H3 methylation (Krogan et al. 2003; Wood et al. 2003). However, many eukaryotes have multiple methyltransferases in addition to Set1 orthologues, and demethylases also contribute to the generation of the H3K4me1/2/3 pattern (Shilatifard 2012; Cenik and Shilatifard 2021; Zhou and Ma 2008; Oya et al 2022). Notably, the targeting mechanisms and significance of H3K4 demethylases for the generation of H3K4me1/2/3 distribution patterns remain largely unexplored.

Genetic screens for factors mediating silencing by H3K9me have identified LYSINE-SPECIFIC DEMETHYLASE 1-LIKE 2 (LDL2) in Arabidopsis (Inagaki et al. 2017). LDL2 is a member of four *Arabidopsis* homologues of human LSD1, a FAD-dependent lysine-specific histone H3K4 demethylase (Shi et al. 2004; Jiang et al. 2007; Rudolph et al. 2007). LDL2 mediates the removal of H3K4me1 from the gene body that accumulates H3K9me2, which in turn directs transcriptional repression. FLOWERING LOCUS D (FLD), another member of the LSD1-like proteins in Arabidopsis, controls transcriptional elongation by removing H3K4me1 from bodies of convergently or bidirectionally transcribed genes (Inagaki et al. 2021). These results suggest that H3K4me1 positively controls transcription and that H3K4me1 demethylases contribute to proper epigenome patterning.

Another LSD1-like protein, LDL3, demethylates H3K4me2 in thousands of protein-coding genes (Ishihara et al. 2019). The *ldl3* mutants compromise shoot regeneration from callus, likely because H3K4me2 demethylation by LDL3 is required for the activation of genes involved in shoot regeneration (Ishihara et al. 2019). In the *ldl3* mutant, the expression level of the flowering repressor gene *FLOWERING LOCUS C* (*FLC*) is decreased, and flowering is accelerated compared to wild type (WT) plants (Martignago et al. 2019). Although these results demonstrate the importance of LDL3-mediated H3K4me2 removal for gene activation, it remains unknown how LDL3 is recruited to its target genes.

To elucidate the chromatin-targeting mechanism of LDL3, we used a machine learning algorithm to identify candidate chromatin features that are targeted by LDL3 for its recruitment. Subsequent genetic and biochemical analyses on the identified candidate features (histone modifications and RNAPII phosphorylation) revealed that LDL3 interacts with CTD-phosphorylated RNAPII and demethylates H3K4me2 cotranscriptionally. Based on these and previous results (Liu et al. 2019; van Dijk et al. 2010; Ishihara et al. 2019; Oya et al. 2022; Wang et al. 2022), we propose that H3K4me2 in plants functions as a memory of the inactive chromatin state, which is established through the interaction between the H3K4me2 demethylase and transcription machinery.

## Results

### Factors associated with LDL3-dependent H3K4me2 removal

To determine genes from which LDL3 removes H3K4me2, we first investigated H3K4me2 levels in shoot tissues in the WT and the *ldl3* mutants by chromatin immunoprecipitation sequencing (ChIP-seq). A total of 7,725 genes with increased H3K4me2 in *ldl3* compared with the WT were defined as LDL3 targets (Fig. 1A). These LDL3 target genes also hyperaccumulate H3K4me2 in the roots and calli of *ldl3* mutants (Ishihara et al. 2019) (Appendix Fig. S1A, B), although the extent of hyperaccumulation varies among tissues.

**Figure 1.**
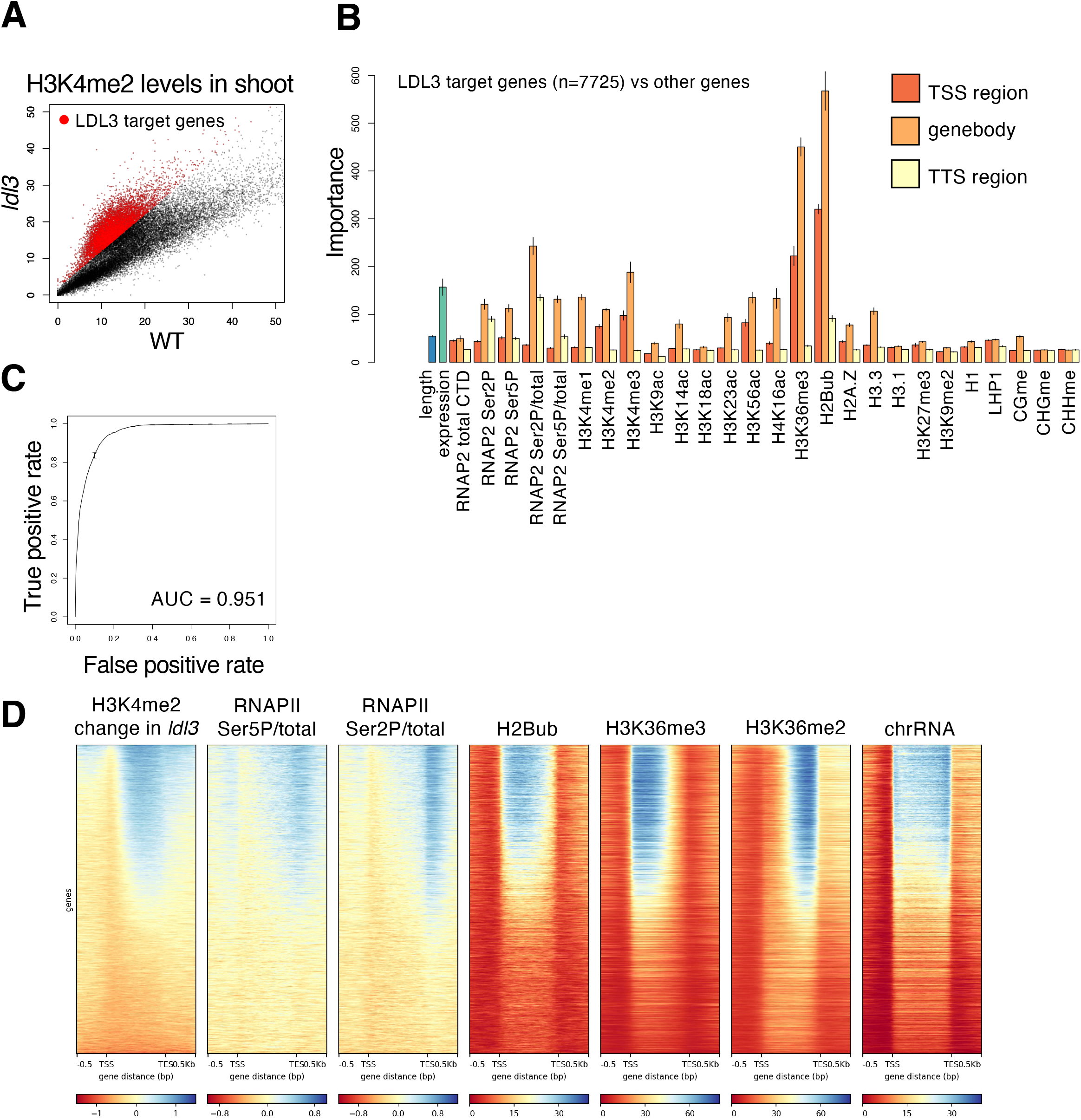
Factors associated with LDL3-dependent H3K4me2 removal. (A) H3K4me2 levels in shoot tissues of the *ldl3* mutants compared with that of WT. Each dot represents RPKM within each protein-coding gene. Red dots, protein-coding genes with hyper H3K4me2 in *ldl3* (H3K4me2 levels in *ldl3* (RPKM) - H3K4me2 levels in *ldl3* (RPKM) > 2; 7,725 genes). (B) Chromatin features predictive of the genes that gain H3K4me2 in the *ldl3* mutant in the random forest models. Error bars represent the standard deviation of the five repeats of training. (C) ROC plot showing the prediction accuracy of the random forest models. (D) Enrichment heatmap depicting ChIP–seq normalized reads. The protein-coding genes were sorted based on increased gene-body H3K4me2 levels in *ldl3*.

We next explored the mechanism by which LDL3 targets specific genes. We reasoned that LDL3 recognizes specific chromatin feature(s), such as epigenome marks and chromatin proteins, and thus, we screened for chromatin feature(s) that distinguish LDL3 demethylation target genes from the other genes using the machine learning algorithm, random forest. The features examined included gene length, expression level, DNA methylation, histone modifications, and RNAPII levels (Fig. 1B). The features that could distinguish LDL3 target genes and others were histone H2B monoubiquitination (H2Bub), H3K36me3, and phosphorylation/total ratio of RNAPII CTD in the gene body (Fig. 1B, C); H2Bub, H3K36me3, and phosphorylated RNAPII highly accumulated in LDL3 target genes compared to the other genes (Fig. 1D).

### Defects in RNAPII phosphorylation and transcriptional elongation mimic loss of LDL3 function

We next genetically tested whether each of the identified candidate factors functions upstream of H3K4me2 demethylation by LDL3. In the *hub1/2* mutant that lacks H2Bub (Cao et al. 2008) (Appendix Fig. S2A) and the *sdg8* mutant that lacks H3K36me3 (Dong et al. 2008; Xu et al. 2008; Oya et al. 2022), H3K4me2 levels were unaffected (Appendix Fig. S2B, C). These results indicate that H2Bub and H3K36me3 are dispensable for LDL3-mediated H3K4me2 removal, although the possibility remains that those features work redundantly in LDL3 recruitment.

To test whether RNAPII phosphorylation acts upstream of H3K4me2 demethylation by LDL3, we examined mutants of an RNAPII CTD kinase, CDKF;1. The *cdkf;1* mutation decreases the levels of both Ser5 and Ser2 phosphorylation (Hajheidari et al. 2012). If RNAPII phosphorylation functions upstream of H3K4me2 demethylation by LDL3, the *cdkf;1* mutation will affect LDL3-induced H3K4me2 demethylation. Indeed, the *cdkf;1* mutant plants showed elevated H3K4me2 levels specifically in genes affected by LDL3 (Fig. 2A, B, C, D, E, Appendix Fig. S3A, B).

**Figure 2.**
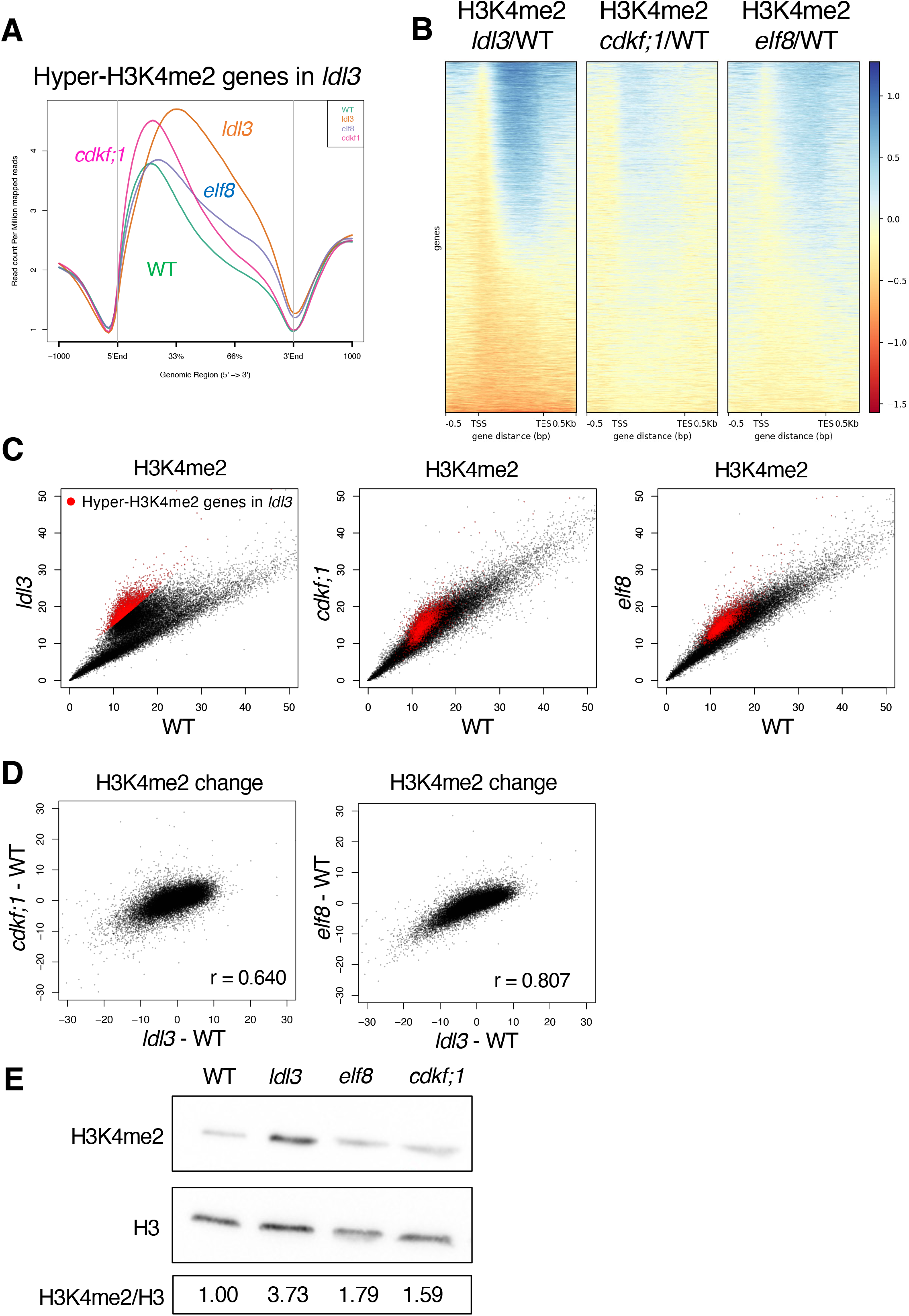
Defects in RNAPII phosphorylation and transcriptional elongation mimic loss of LDL3 function. (A) Averaged profiles of H3K4me2 around LDL3 target genes in each genotype (WT, *ldl3, elf8, cdkf;1*). The numbers of genes analyzed are 7367. The ribbons indicate s.e.m. (B) The *elf8* and *cdkf;1* mutant plants show an increase in H3K4me2 for genes affected by *ldl3*. Heatmap of changes in H3K4me2 levels is shown in the three mutants. The protein-coding genes were sorted based on the increase in gene-body H3K4me2 levels in *ldl3*. (C) H3K4me2 levels in each of the mutants compared with WT. Each dot represents RPKM within each protein-coding gene. Red dots, protein-coding genes with hyper H3K4me2 in *ldl3* (H3K4me2 levels in *ldl3* (RPKM) - H3K4me2 levels in *ldl3* (RPKM)>5; 2,809 genes). (D) Relationships between changes in H3K4me2 levels (RPKM) in *elf8* and *cdkf;1*, and in *ldl3* within each of protein-coding genes compared to WT. (E) Western blotting of H3K4me2 on bulk histone extracted from the mutants. The ratios (H3K4me2/H3) of quantified western blotting signals are shown as the relative value to WT.

Ser5 and Ser2 phosphorylation of the RNAPII CTD are implicated in transcriptional elongation (Harlen and Churchman 2017). We therefore explored the relationship between transcriptional elongation and LDL3-mediated H3K4me2 demethylation using mutants of the transcriptional elongation factor Paf1 complex (Paf1C). Paf1C is known to be involved in RNAPII CTD phosphorylation in yeast and animals (Dronamraju and Strahl 2014; Yu et al. 2015). Paf1C has also been reported to colocalize with phosphorylated RNAPII in plants (Antosz et al. 2017). *Arabidopsis* Paf1C consists of VERNALIZATION INDE-PENDENCE2 (VIP2)/EARLY FLOWERING7 (ELF7), VIP4, VIP5, VIP6/ELF8, and CDC73/PLANT HOMOLOGOUS TO PARAFIBROMIN (PHP) (Oh et al. 2004; Yu and Michaels 2010; Park et al. 2010). The mutants of these Paf1C component genes showed elevated H3K4me2 levels compared with WT, especially in the LDL3 target genes (Appendix Fig. S4). This result suggests that Paf1C is required for H3K4me2 demethylation in the LDL3 target genes. In subsequent analyses, we further investigated the function of Paf1C on H3K4me2 demethylation using the *elf8* mutant as a representative of the Paf1C components. The H3K4me2 elevation occurred in largely overlapping genes in the *elf8, ldl3* and *cdkf;1* mutants. (Fig. 2B, C, D Appendix Fig. S3B). Previous studies (Oh et al. 2004) proposed that Paf1C did not affect global H3K4me2/me3 based on western blotting results, but our ChIP-seq results showed that Paf1C significantly affects H3K4me2 distribution. H2Bub and H3K36me2/3, which random forest analysis also identified, were altered in the *elf8* mutant (Appendix Fig. S5A, B, C) (Oh et al. 2008; Liu et al. 2019), but these modifications were not altered in the *ldl3* mutant, suggesting that they may be regulated by ELF8/Paf1C, independently of LDL3. In fact, Paf1C has been reported to bind the H3K36 methyltransferase SDG8 (Yang et al. 2016).

In the *ldl3* mutant, increased H3K4me2 was associated with a decrease in H3K4me1, likely because H3K4me1 is the product of H3K4me2 demethylation by LDL3 (Ishihara et al. 2019). Both the *cdkf;1* and *elf8* mutants showed changes in the H3K4me1 pattern analogous to that of *ldl3* (Appendix Fig. S6A, B, C). These results indicate that transcriptional elongation and/or RNAPII phosphorylation functions upstream of H3K4me2 demethylation by LDL3.

LDL3 expression levels were not reduced in the *cdkf;1* or in the *elf8* mutant (Appendix Fig. S7), indicating that Paf1C and CDKF;1 likely affect LDL3 function or recruitment, not its transcription. In addition, the *cdkf;1* and *elf8* mutants compromised shoot regeneration from callus similar to the *ldl3* mutant (Ishihara et al. 2019) (Appendix Fig. S8A, B). These results suggest a novel pathway in which LDL3-mediated H3K4me2 removal is controlled through the interaction between LDL3 and elongating RNAPII, and this pathway is indispensable for regeneration.

### LDL3 binds to phosphorylated RNAPII in vivo

To test whether the effect of RNAPII on H3K4me2 is mediated by its direct binding to LDL3, we performed coimmunoprecipitation (Co-IP) analysis using transgenic plants that expressed FLAG-tagged LDL3. RNAPII was pulled down by IP with an antibody against FLAG (Fig. 3A), which showed that LDL3 interacts with RNAPII. LDL3 bound much more strongly to CTD-phosphorylated RNAPII than to unphosphorylated RNAPII (Fig. 3A). Taken together with the colocalization of phosphorylated RNAPII and LDL3 targets shown in the heatmap (Fig. 1D), it is likely that LDL3 binds to phosphorylated RNAPII and removes H3K4me2 cotranscriptionally. Interestingly, the results showed that the other LSD1 family H3K4 demethylases, FLD and LDL2 did not bind to RNAPII (Fig. 3B). Among the three structurally related H3K4 demethylases, binding to RNAPII was unique to LDL3.

**Figure 3.**
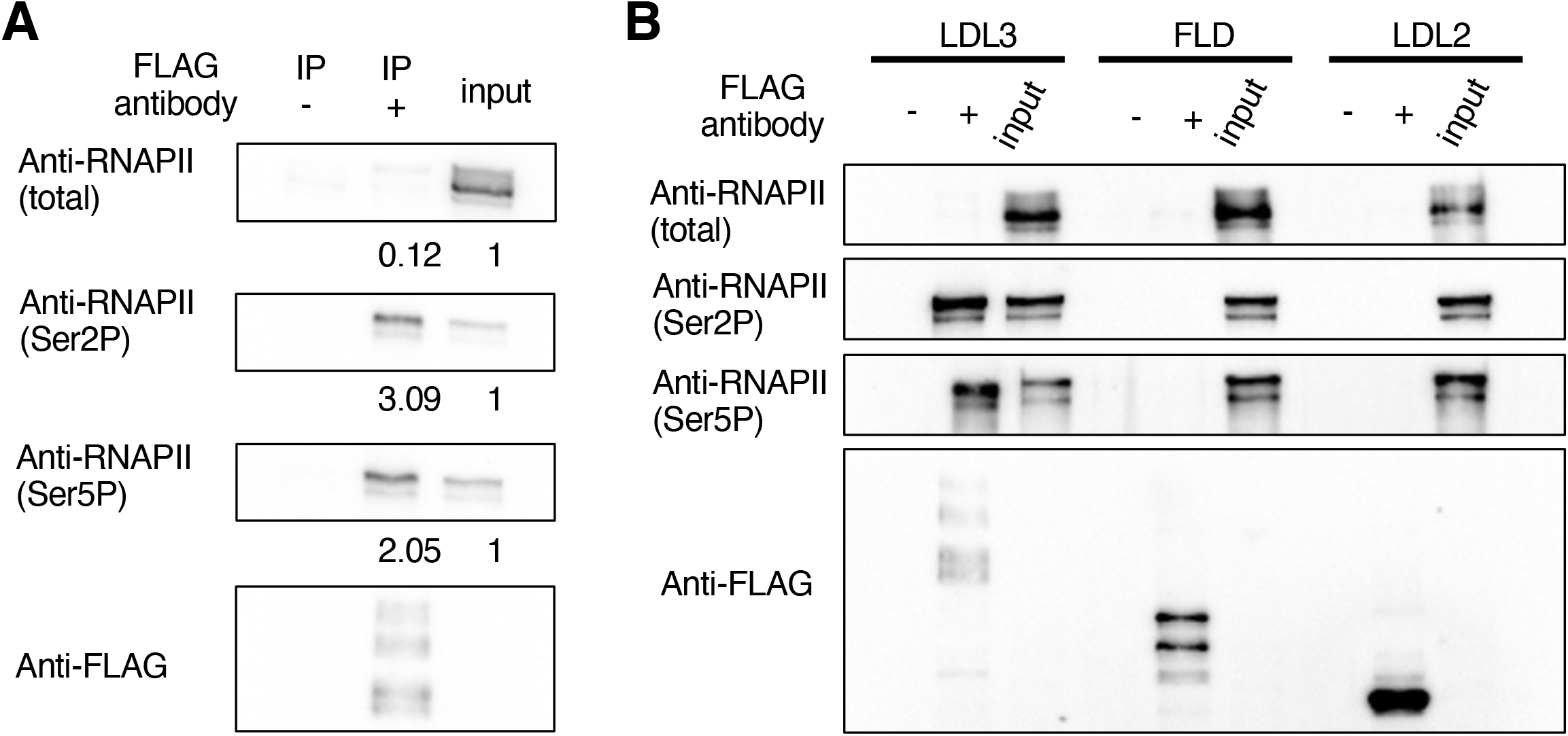
LDL3 is bound to phosphorylated RNAPII in vivo. (A) LDL3 was immunoprecipitated with anti-FLAG beads (IP: FLAG) from whole cell lysates and analyzed by immunoblotting using antibodies against Ser5P, Ser2P or total RNAPII CTD. The ratios (IP/input) of quantified western blotting signals are shown at the bottom of each blot. (B) Co-IP experiments testing the binding between RNAPII and each of the LSD1 family proteins (LDL3, FLD, LDL2).

### RNAPII phosphorylation and transcriptional elongation are differentially altered in *elf8* and *cdkf;1* mutants

The results above suggest that phosphorylated RNAPII mediates the function of LDL3 in removing H3K4me2. We therefore examined the levels of phosphorylated RNAPII in the *cdkf;1* and *elf8* mutants by western blotting. As expected from its coding protein, the *cdkf;1* mutant showed decreases in RNAPII phosphorylation levels (both Ser2P and Ser5P) (Fig. 4A), which was consistent with a previous study (Hajheidari et al. 2012). Unexpectedly, however, the proportion of phosphorylated RNAPII was increased in *elf8* (Fig. 4A). To further clarify the effect of *cdkf;1* and *elf8* on RNAPII phosphorylation, we examined genome-wide patterns of RNAPII phosphorylation by ChIP-seq analysis. Consistent with the idea that the loss of phosphorylation in the *cdkf;1* mutant mediates its effect on LDL3 function, the ratio of Ser2P/total RNAPII in the gene body and around the transcription termination site (TTS) was decreased in *cdkf;1* in the LDL3 target genes (Fig. 4B, C).

**Figure 4.**
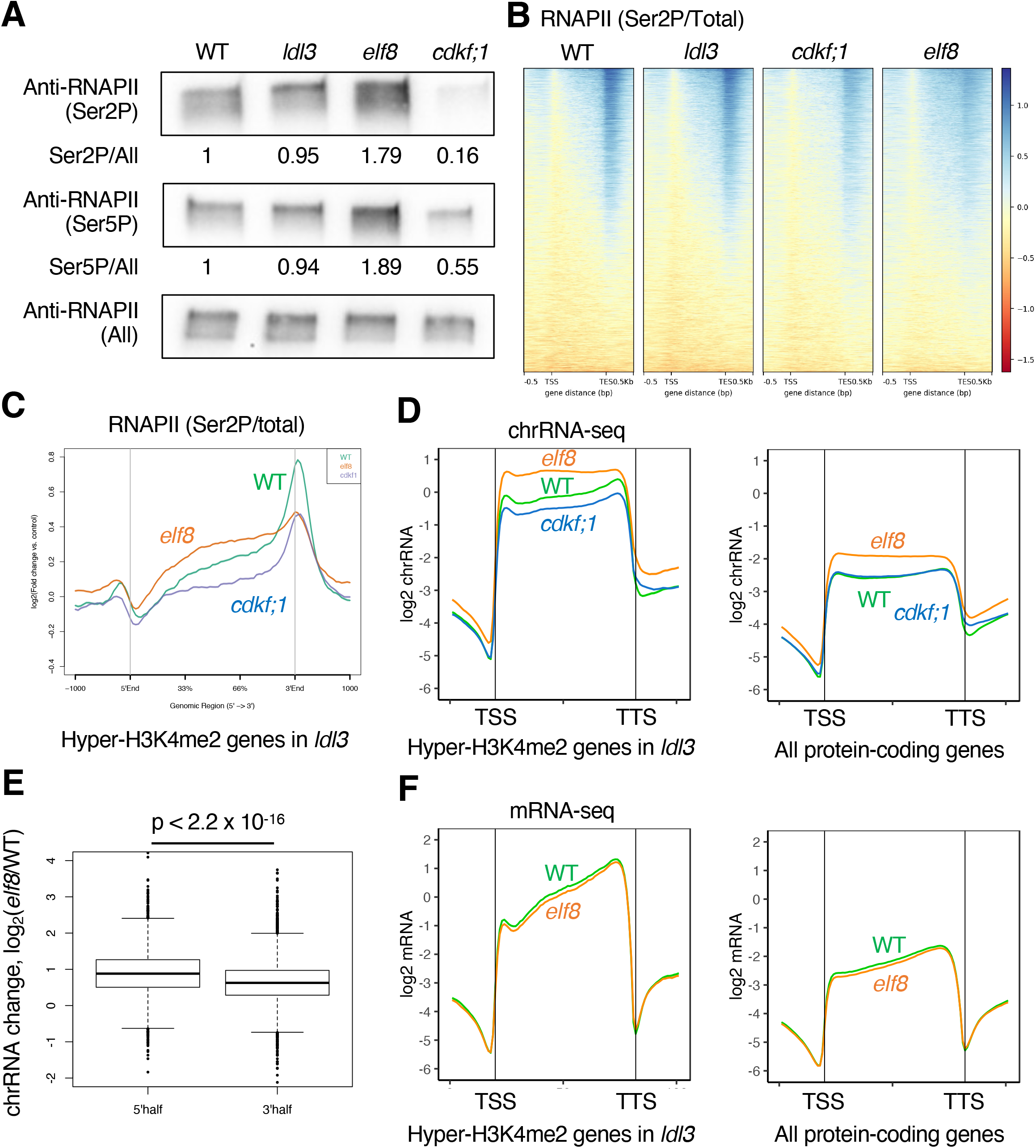
RNAPII phosphorylation and transcriptional elongation are differentially altered in *elf8* and *cdkf;1* mutants. (A) Western blotting of protein extracted from seedlings of each genotype. Membranes were probed with antibodies against Ser5P or Ser2P or total RNAPII CTD. The ratios (phosphorylated/total) of quantified western blotting signal are shown as the relative value to WT. (B) Heatmaps of RNAPII Ser2P levels (Ser2P/total CTD ratio) are shown in WT and the three mutants. The protein-coding genes were sorted based on increased gene-body H3K4me2 levels in *ldl3*. (C) Averaged profiles of Ser2P levels around the LDL3 target genes for each genotype (WT, *elf8, cdkf;1*). The numbers of genes analyzed are 7,367. The ribbons indicate s.e.m. (D) Averaged profiles of chrRNA around LDL3 target genes (n=7,367) and all protein-coding genes (n=27,206) for each genotype (WT, *elf8, cdkf;1*). The ribbons indicate s.e.m. (E) Box plots comparing the ratio of chrRNA levels in 5’ half of the gene body (left) and 3’ half of the gene body (right) between the WT and *elf8*, in LDL3 target genes (7,367 genes). The P values are based on Welch’s two-sample t-test. (F) Averaged profiles of mRNA around LDL3 target genes (n=7,367) and all protein-coding genes(n=27,206) for each genotype in WT and *elf8*. The ribbons indicate s.e.m.

In contrast, the ratio of Ser2P/total RNAPII was increased in the gene body in the *elf8* mutant, and the pattern was skewed towards the upstream region (Fig. 4B, C). The ratio of Ser5P/total RNAPII in the gene body and around TTS also increased in *elf8* and decreased in *cdkf;1* (Appendix Fig. S9A, B). These results suggest that although phosphorylation of RNAPII is necessary for LDL3 function, *elf8* mutation affects LDL3 function through a different pathway.

To determine how the loss of Paf1C and CDKF;1 affects transcription dynamics and LDL3 function, we conducted chromatin-bound RNA sequencing (chrRNA-seq) to examine nascent transcribing RNAs (Nojima et al. 2015; Inagaki et al. 2021). The *elf8* mutant showed a remarkable change in transcriptional dynamics, with a large amount of nascent RNA detected near the TSS (Fig. 4D). This trend was more pronounced in the LDL3 target genes. In contrast, chrRNA levels were slightly reduced in *cdkf;1* (Fig. 4D). These chrRNA-seq results were consistent with the RNAPII ChIP-seq results. In the *elf8* mutant, RNAPII phosphorylation levels and chrRNA levels were increased in the gene body, especially in the 5’ half of the gene (Fig. 4E). Since mRNA levels were not increased (Fig. 4F), transcriptional elongation was considered to be retarded. Conversely, in the *cdkf;1* mutant, RNAPII phosphorylation and chrRNA levels were reduced, suggesting an overall reduced transcriptional activity caused by reduced RNAPII phosphorylation. The patterns of Ser2/total RNAPII and chrRNA were not largely affected in the *ldl3* mutant (Appendix Fig. S9C, D), suggesting that the increased H3K4me2 level is a consequence but not a cause of altered RNAPII dynamics and phosphorylation. Taken together, Paf1C and CDKF;1 regulate RNAPII phosphorylation and transcriptional elongation in different ways.

H3K4me2 profiles also differed between *elf8* and *cdkf;1*. In the *elf8* mutant, the H3K4me2 level increased in the 3’ half of the target genes (Fig. 2A, Appendix Fig. S3A), which was consistent with the stalled transcription in the 5’ half of the gene body (Fig. 4D, E). In the *cdkf;1* mutant, H3K4me2 increased while maintaining the WT profile (Fig. 2A, Appendix Fig. S3A), possibly reflecting reduced overall transcription activity (Fig. 4D). Differential changes in RNAPII phosphorylation and transcriptional elongation in *cdkf;1* and *elf8* may generate different patterns of H3K4me2 increase.

### The *elf8* mutation affects H3K4me2 through retardation of transcription in cis

We further explored the mechanism by which Paf1C regulates LDL3-mediated H3K4me2 demethylation. The direct interaction between LDL3 and phospho-RNAPII (Fig. 3A) supports the hypothesis that Paf1-mediated transcriptional elongation directly facilitates the local function of LDL3, but it is still possible that change(s) in the expression of other gene(s) indirectly affect LDL3 function. We reasoned that such indirect effects would cause elevation of H3K4me2 irrespective of the local effect of Paf1 on transcriptional elongation. Thus, we tested whether the *elf8*-induced retardation of transcriptional elongation in each gene correlates with its effect on H3K4me2. Indeed, we found a clear positive correlation between estimated transcriptional retardation and H3K4me2 accumulation in the *elf8* mutant (Fig. 5A, B, D). In WT, ELF8 localized to these genes that showed hyperaccumulation of chrRNA, total and phosphorylated RNAPII, and H3K4me2 in *elf8* (Fig. 5C, D). These results suggest that RNAPII-associated Paf1C ensures transcriptional elongation and demethylation of H3K4me2, the latter of which is most likely through LDL3 function.

**Figure 5.**
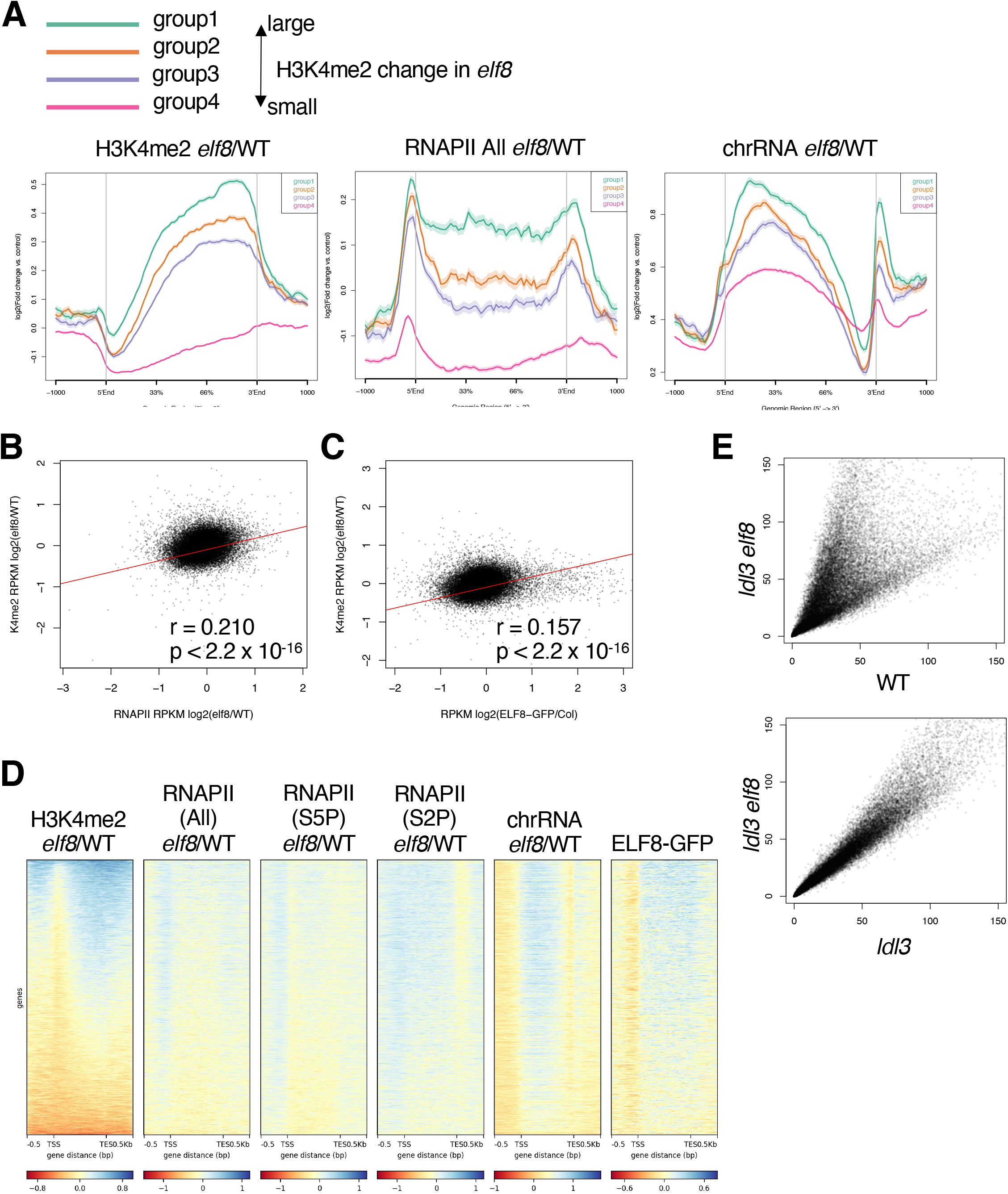
The *elf8* mutation affects H3K4me2 through retardation of transcription in cis. (A) Averaged profiles of H3K4me2 changes, chrRNA changes, and RNAPII changes between WT and *elf8*. All protein-coding genes were grouped based on levels of change in H3K4me2. Group1: 5 < *elf8*-WT (RPKM) (green, 2,809 genes); Group2: 3 < *elf8*-WT (RPKM) < 5 (orange, 2,919 genes); Group3: 1 < *elf8*-WT (RPKM) < 3 (blue, 3234 genes); and Group4: *elf8*-WT (RPKM) < 1 (pink, 17,721 genes). (B) Relationships between changes in RNAPII levels (RPKM) and changes in H3K4me2 levels (RPKM) in *elf8* within each protein-coding gene compared to WT. (C) Relationships between the ELF8 localization corrected with a non-transgenic control and changes in H3K4me2 levels (RPKM) in *elf8*. (D) Heatmaps depicting changes in H3K4me2 levels, RNAPII levels (total or phosphorylated), and chrRNA levels in the *elf8* mutants, and the ELF8 localization. ELF8 localization is corrected with a nontransgenic control. (E) H3K4me2 levels in *ldl3 elf8* double mutant compared with WT (top) or (bottom). Each dot represents the RPM within each protein-coding gene. Developmental phenotypes of the double mutant and each of the single mutants are shown in Appendix Fig. S10B.

We also genetically tested whether Paf1C and LDL3 work through the same pathway for H3K4me2 regulation by analysing the *elf8 ldl3* double mutant. The *elf8 ldl3* double mutant showed an increase in H3K4me2 similar, although not identical, to that in the *ldl3* single mutant (Fig. 5E, Appendix Fig. S10A), suggesting that the accumulation of H3K4me2 in the *elf8* mutant is mostly through compromising the function of LDL3.

### LDL3 disrupts the positive correlation between H3K4me2 and transcription

In WT, the H3K4me2 level was negatively correlated with the transcription level (Fig. 6A), which was consistent with previous studies (Liu et al. 2019). Interestingly, however, we observed a positive correlation between H3K4me2 and transcription levels in the *ldl3* mutant (Fig. 6A). The negative correlation was also attenuated in the *cdkf;1* and *elf8* mutants, although the extent was weaker than that in *ldl3* (Appendix Fig. S11). These results suggest that LDL3 disrupts the positive correlation of transcription and H3K4me2 by removing H3K4me2 from transcribed genes. In other words, LDL3 demethylates H3K4me2 cotranscriptionally to establish the negative correlation between transcription and H3K4me2. Furthermore, the positive correlation between transcription and H3K4me1 levels, which was seen in WT, was attenuated in the *ldl3, cdkf;1*, and *elf8* mutants (Fig. 6A, Appendix Fig. S11), suggesting that conversion from H3K4me2 to H3K4me1 by LDL3 significantly contributes to generating the positive correlation between transcription and H3K4me1. These results indicate that LDL3 demethylates H3K4me2 cotranscriptionally to establish higher H3K4me1 and lower H3K4me2 levels within actively transcribed genes.

**Figure 6.**
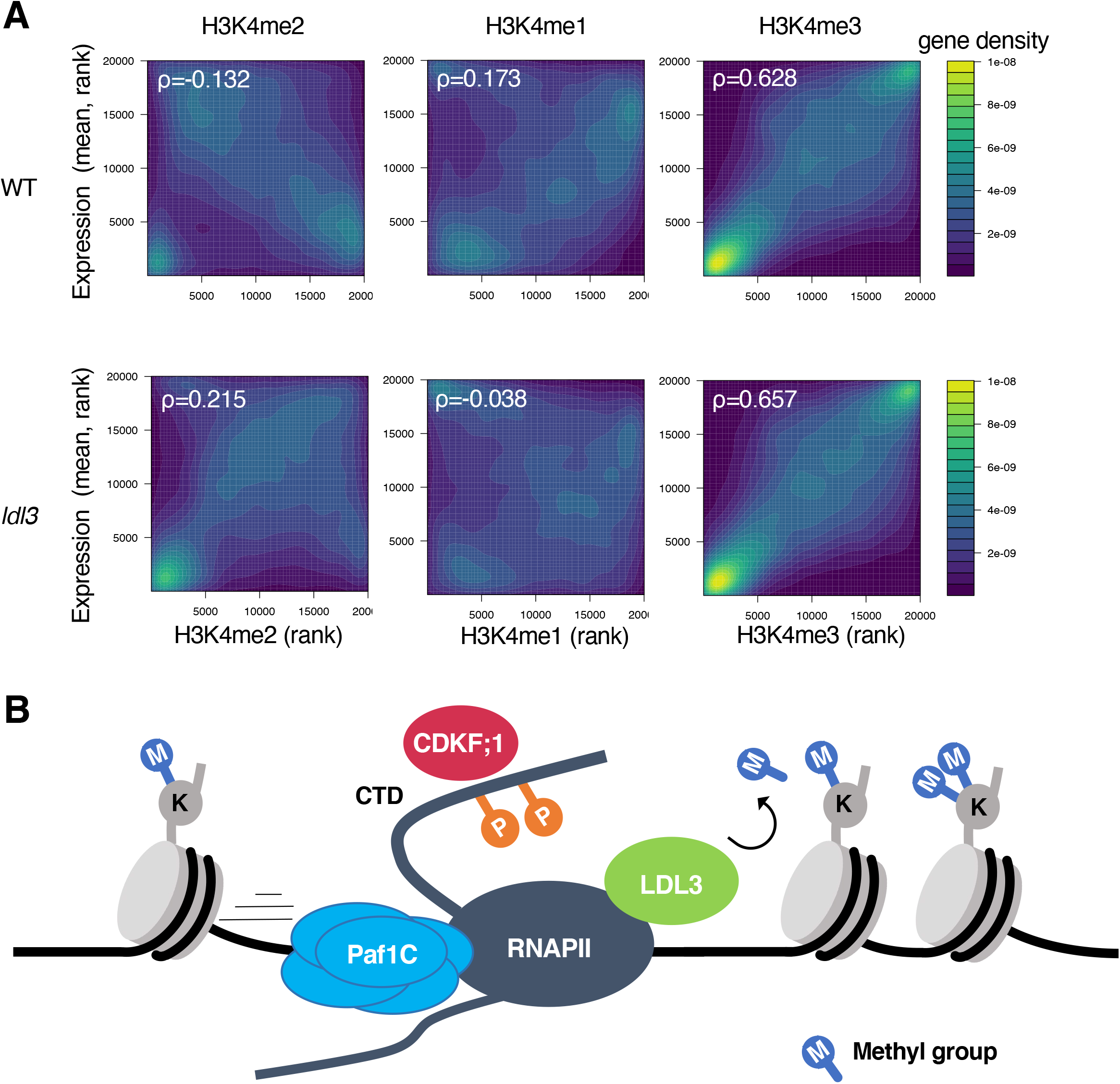
*LDL3* disrupts the positive correlation between H3K4me2 and transcription. (A) Correlations between H3K4me and transcription level of protein-coding genes in wild-type (WT) and *ldl3* mutant plants. Protein-coding genes are ranked by H3K4me1 or me2 or me3 (x-axis, RPKM) and expression levels (y-axis, chrRNA-seq, RPKM). The densities of genes are visualized as heat maps. ρ: Spearman’s correlation coefficient. (B) Model for co-transcriptional H3K4me2 demethylation by LDL3. LDL3 recognizes RNAPII phosphorylation and binds to RNAPII, and thus H3K4me2 is removed co-transcriptionally.

## Discussion

In this study, we searched for candidate factors that determine the functional pathway of LDL3 through an initial screening with random forest analyses and subsequent genetic tests. By this approach, we identified RNAPII phosphorylation and transcriptional elongation as factors functioning upstream of LDL3-mediated H3K4me2 demethylation.

The involvement of RNAPII phosphorylation in LDL3 function was further substantiated by the biochemical results showing that LDL3 binds to phosphorylated RNAPII. Interestingly, such binding was not detected for two other LSD1-type *Arabidopsis* demethylases, LDL2 and FLD, which remove H3K4me1. LDL3, but not LDL2 or FLD, has the SRI domain, which was identified in yeast Set2 (Kizer et al. 2005) as a domain specifically binding to the phosphorylated CTD of RNAPII (Rebehmed et al. 2014). While mammalian LSD1 family proteins do not have the SRI domain, the SRI domain is found in LDL3 orthologues in rice, poplar, the seedless ancient vascular plant *Selaginella moellendorffi*, and even the moss *Physcomitrella patens* (Rebehmed et al. 2014) (Appendix Fig. S12), suggesting that cotranscriptional H3K4 demethylation via the SRI domain may be conserved among land plants.

The *elf8* mutant showed an increase in H3K4me2 in the LDL3 targets, most likely through retardation in transcriptional elongation. Interestingly, however, that was associated with an increase, rather than a decrease, in RNAPII phosphorylation. On the other hand, a delay in transcriptional elongation was prominent in the *elf8* mutant but not in the *cdkf;1* mutant. These results suggest that *elf8* and *cdkf;1* affect LDL3 function through distinct pathways. Mutations of the other Paf1C components also affect H3K4me2, suggesting that their function as a Paf1 complex is important for H3K4me2 demethylation. The H3K4me2 pattern in the *elf8 ldl3* double mutants also suggests that the effect of Paf1C on H3K4me2 is mediated by LDL3. The *elf8* mutation affects H3K4me2 in the genes that are bound by ELF8 and shows retardation of transcriptional elongation in *elf8*. Taken together, these results demonstrate that defects in Paf1C lead to transcriptional retardation and affect LDL3-mediated H3K4me2 demethylation directly.

Cotranscriptional H3K4 methylation has been reported in yeast and animals (Woo et al. 2017). RNAPII phosphorylation is known to stimulate H3K4 methylation by Set1-COMPASS, and Paf1C is also important for Set1 recruitment (Krogan et al. 2003; Ng et al. 2003). The cotranscriptional effect is also observed in the Set1-type *Arabidopsis* methyltransferases ATXR3 and ATXR7, which deposit H3K4me3 and H3K4me1, respectively (Oya et al. 2022), although Paf1C is not essential in this mechanism in *Arabidopsis* (Oh et al. 2004). On the other hand, our study shows that Paf1C and phosphorylation of RNAP II contribute to the LDL3-mediated removal of H3K4me2 in *Arabidopsis*. LDL3 establishes the plant-specific negative correlation between H3K4me2 and transcription by the transcription-driven demethylation of H3K4.

Thus, transcription induces high H3K4me1/3 states by Set1-type methyltransferases, but it also induces low H3K4me2 by transcription-coupled removal of H3K4me2 (Fig. 6B). As it has been proposed in previous studies that transcription-coupled H3K4 regulation may establish a memory of recent transcriptional activity (Ng et al. 2003; Ding et al. 2012; Muramoto et al. 2010), it is possible that H3K4me2 removal also functions as a memory of transcription. This hypothesis is consistent with reports that H3K4me2 is a repressive modification in plants (*Arabidopsis* and rice) (Liu et al. 2019; Wang et al. 2022). It is also known that H3K4me2 levels are decreased in transcriptionally upregulated genes under dehydration stress (van Dijk et al. 2010). The *ldl3* mutant showed decreased *FLC* expression levels, which affects the developmental transition to environmental stimuli (Martignago et al. 2019). Furthermore, LDL3 function is necessary for shoot regeneration from callus by “priming” the expression of key factors (Ishihara et al. 2019). Based on these previous studies and current results, we propose that active demethylation by LDL3, driven by transcriptional elongation, functions as a memory to control developmental plasticity and robust gene control in plants. This concept could be further extended in the future using *Arabidopsis* histone MTase mutants specifically affecting H3K4me2 (Oya et al. 2022). Finally, binding to the elongating transcription machinery has also been reported in the human LSD1 homologue, LSD2, which functions specifically for H3K4me2 (Fang et al. 2010). It would be interesting to determine whether a similar strategy has convergently evolved to control diverse functions of H3K4me2 in animals and plants.

## Materials and methods

### Plant materials and growth condition

*Arabidopsis thaliana* strain Columbia-0 (Col-0) was used as wild type (WT). The mutant alleles used in this study were: *ldl3-1* (GABI_092C03) (Ishihara et al, 2019), *elf7-2* (SALK_046605) (Tamada et al, 2009), *elf7-3* (SALK_019433) (Cao et al, 2008; Li et al, 2019), *elf8-1* (SALK_090130), *elf8-2* (SALK_065364) (Shiraya et al, 2008), *vip3-1* (SALK_139885) (Fal et al, 2017, 2019), *vip4-4* (SALK_039374) (He et al, 2004; Li et al, 2019), *vip5-1* (SALK_062223) (Oh et al, 2004), *cdc73-1* (SALK_150644) (Yu & Michaels, 2010), *cdkf;1-1* (SALK_148336) (Hajheidari et al, 2012), *hub1-4* (SALK_122512) (Cao et al, 2008), *hub2-2* (SALK_071289) (Cao et al, 2008), *sdg8* (*ashh2-1*, SALK065480) (Grini et al, 2009), all of which are in the Col-0 background. Plants were grown on Murashige and Skoog (MS) media supplemented with 1% sucrose and solidified with Bacto agar under long day (16 h light/8 h darkness) photoperiods at 22ºC.

To make the pLDL3::LDL3-3xFLAG fusion construct, an LDL3 genomic DNA fragment (2.5 kb upstream of ATG to just before the stop codon) followed by 3xFLAG sequence was cloned into the pGreenII 0179 vector. The plasmid was transformed into the *ldl3-1* mutant via Agrobacterium tumefaciens GV3101::pMP90. In the same way, the pELF8::ELF8-GFP fusion construct was cloned from an ELF8 genomic DNA fragment (from ∼1 kb upstream of the ATG to just before the stop codon). The plasmid was transformed into the *elf8-2* mutant. FLAG-tagged LDL2 and FLD plants used in this work were described before (Inagaki et al, 2017, 2021).

### ChIP-seq

Chromatin immunoprecipitation sequencing (ChIP-seq) was carried out following (Inagaki et al, 2017) with modifications. 0.5∼1.0 g of aerial parts from 14-day-old seedlings were used as each ChIP sample. Samples were ground in liquid nitrogen and crosslinked for 10 minutes at room temperature in Nuclei isolation buffer (10 mM HEPES pH 7.5, 1 M sucrose, 5 mM KCl, 5 mM MgCl2, 5 mM EDTA, 0.1 % β-mercaptoethanol, 0.6 % Triton X-100, supplemented with 1 tablet/50 ml cOmplete proteinase inhibitor and 1 mM Pefabloc SC (Roche)) supplemented with 1 % formaldehyde. In the case of the LDL3-3xFLAG ChIP shown in Figure 5, the crosslink buffer additionally contains 1.5 mM ethylene glycol bis(succinimidyl succinate) (EGS). In the case of RNAPII ChIP, the formaldehyde concentration was 2%. The crosslinking reaction was stopped with 130 mM glycine. The suspension was filtered through a 40 μm nylon cell strainer and pelleted by centrifugation at 3,000 rpm at 4 °C for 10 min. The pellet was resuspended in 300 μl of nuclei isolation buffer and layered on top of 500 μl of nuclei separation buffer (10 mM HEPES pH 7.6, 1 M sucrose, 5 mM KCl, 5 mM MgCl2, 5 mM EDTA pH 8.0, 15 % Percoll) and pelleted by centrifugation at 16,000 × g for 5 min at 4 °C. The nuclear pellet was resuspended in 950 μl RIPA without Triton buffer (50mM Tris-HCl pH7.8, 150mM NaCl, 1mM EDTA, 0.1% SDS, 0.1% Sodium deoxycholate and cOmplete proteinase inhibitor).

Sonication was conducted using Covaris S2 Focused-ultrasonicator (Covaris) and milliTUBE 1 ml AFA Fiber (Covaris) with following settings: power mode, frequency sweeping; time, 18-20 min; duty factor, 5%; Cycles per Burst, 200; temperature (water bath), 4–6 °C; and Peak Incident Power, 140 for histones and 105 for epitope-tagged proteins. In the case of RNAPII ChIP, chromatin was sheared using a Picoruptor (Diagenode) (6 cycles of 30 seconds on and 30 seconds off).

The sonicated chromatin was then centrifuged at 13,000 g for 3 min, and the supernatant was added with 50 μL of 20 % Triton X-100 and aliquoted. The chromatin solution was incubated with 1-2 μg of antibodies overnight at 4 ºC. Antibodies used are H3K4me1 (ab8895; Abcam), H3K4me2 (ab32356; Abcam), H3K4me3 (ab8580; Abcam), and anti-H3 (ab1791; Abcam), H3K36me2 (MABI0332; MBL), H3K36me3 (MABI0333; MBL), H2Bub (MM-0029; Medimabs), RNAP2 total CTD (Diagenode, C15100055), RNAP2 phospho S2 (MABI0602; MBL), RNAP2 phospho S5 (MABI0603; MBL), FLAG (F1804; SIGMA).

Then the antibody-chromatin mix was incubated for 2 h at 4 °C with magnetic beads; Dynabeads M280 Sheep anti-mouse IgG in the cases of MBL antibodies, and with Dynabeads Protein G in the cases for the others. Wash, reverse cross-linking, and DNA extraction were conducted as described previously (Inagaki et al, 2021). For epitope-tagged proteins, instead of LiCl buffer (1% IGEPAL, 1% Sodium deoxycholate), half LiCl buffer (0.5% IGEPAL, 0.5% Sodium deoxycholate) was used. Collected DNA was quantified with the Qubit dsDNA High Sensitivity Assay kit (Thermo Fisher Scientific), and 1–2 ng DNA was used to make libraries for Illumina sequencing. The library was constructed with the KAPA Hyper Prep Kit for Illumina (KAPA Biosystems), and dual size-selected using SPRI select beads (Beckman Coulter) to enrich 300–500-bp fragments. The libraries were 50-bp single-end sequenced by HiSeq4000 sequencer (Illumina) inVincent J. Coates Genomics Sequencing Laboratory at UC Berkeley, or 150 bp paired-end sequenced by the HiSeqX Ten sequencer (illumina). Biological replicates were conducted on independently grown plants.

### eChIP-seq

In the experiments shown in Fig. 5E and Appendix Fig. S4, the eChIP method (Zhao et al, 2020), which can be performed with a small amount of plant material, was performed because many mutants that were used in the experiment showed sterility and it was difficult to collect a large amount of material. The results were comparable to those of conventional methods.

0.01∼0.1 g of aerial parts from 14-day-old seedlings were ground in liquid nitrogen and added to Fixation Buffer (phosphate-buffered saline (PBS) with 1% formaldehyde, 1mM Pefabloc SC (Roche) and cOmplete protease inhibitor cocktail (Roche))and incubated at room temperature for 10 min. Glycine was added to a concentration of 0.2 M and further incubated at room temperature for 5 min, after which the supernatant was removed by centrifugation. Pellets were lysed in 180 μL of Buffer S (50 mM HEPES-KOH (pH 7.5), 150 mM NaCl, 1 mM EDTA, 1 % Triton X-100, 0.1 % sodium deoxycholate, 1 % SDS) for 10 min at 4 ºC. The homogenate was mixed with 720 μL of Buffer F (50 mM HEPES-KOH (pH 7.5), 150 mM NaCl, 1 mM EDTA, 1 % TritonX-100, 0.1 % sodium deoxycholate). The chromatin was fragmented into 200 ∼ 600 bp by sonication using Picoruptor (diagenode). The rest of the procedure was the same as in (Zhao et al, 2020). The antibodies used were H3: ab1791 (Abcam) and H3K4me2: ab7766 (Abcam).

### mRNA-seq

Total RNA was isolated from the aerial part of one 14-day-old seedling grown on MS media, using the RNeasy Plant Mini Kit (Qiagen), and treated with DNase I (Takara). Libraries for mRNA-seq were constructed using the KAPA Stranded RNA-seq Library Preparation Kit according to the manufacturer’s instructions. First, poly(A) RNA was purified and then fragmented by heating purified RNA at 94 °C for 7 min in the fragment, prime, and elute buffer. Then, double-strand cDNA was synthesized, and adapter-ligated libraries were constructed. Two independent biological replicates were analyzed for each genotype.

### mRNA-seq

Total RNA was isolated from the aerial part of one 14-day-old seedling grown on MS media, using the RNeasy Plant Mini Kit (Qiagen), and treated with DNase I (Takara). Libraries for mRNA-seq were constructed using the KAPA Stranded RNA-seq Library Preparation Kit according to the manufacturer’s instructions. First, poly(A) RNA was purified and then fragmented by heating purified RNA at 94 °C for 7 min in the fragment, prime, and elute buffer. Then, double-strand cDNA was synthesized, and adapter-ligated libraries were constructed. Two independent biological replicates were analyzed for each genotype.

### chrRNA-seq

First, nuclei were extracted from approximately 0.5 g of aerial parts from 14-day-old seedlings grown on MS media following the protocol as described previously (Inagaki et al, 2021) with the following modifications. The isolated nuclei were resuspended in 500 μl of NUN1 buffer (20 mM Tris–HCl, pH 8.0, 75 mM NaCl, 0.5 mM EDTA, 50 % glycerol, and 1x cOmplete EDTA-free Protease Inhibitor Cocktail) followed by 500 μl of NUN2 buffer (20 mM HEPES-KOH pH 7.6, 7.5 mM MgCl2, 0.2 mM EDTA, 300 mM NaCl, 1 M urea, 1 % NP40, cOmplete EDTA-free Protease Inhibitor Cocktail). The solution was incubated at 4 ºC for 15 min, with vortexing every 3 min. The chromatin pellet was precipitated by performing 15,000 g centrifuge at 4 ºC for 3 min. The chromatin pellet was resuspended in 100 μl of lysis buffer (10 mM Tris–HCl, pH 7.8, 10 mM EDTA, 0.5 % SDS). These samples were used for chrRNA-seq and RNAPII western blotting. chrRNA was extracted using miRNeasy kit (Qiagen). Ribosomal RNA was depleted and the remaining DNA was depleted using the KAPA RiboErase kit. The chrRNA-seq libraries were constructed from 50 ng of purified RNA using the KAPA Stranded mRNA-seq Kit, skipping the poly-A selection step and performing the RNA fragmentation step at 94 ºC for 5 min. Two biological replicates were analyzed. 150-bp paired-read sequences were obtained using the HiSeq X Ten sequencer (Illumina) at Macrogen Japan Corp.

### Data analysis

The mRNA-seq, ChIP–seq, and chrRNA-seq data were processed as described in (Inagaki et al, 2021).

For the random forest analysis, each gene was divided into the TSS region, the TTS region and the gene body. The TSS (or TTS) region here was defined as 200 bp upstream and downstream from the TSS (or TTS) combined, and the gene body was defined as the region spanning the TSS to the TTS. Short genes (<400 bp) were omitted. The objective variable was defined as follows: Hyper-H3K4me2 genes in the *ldl3* mutant are genes with [RPKM of H3K4me2 in *ldl3*] − [RPKM of H3K4me2 in *ldl3*] > 2, which are defined as ‘LDL3 target genes’ (n = 7,725), and the others are ‘non-target’.

The numbers of LDL3 target and non-target genes were balanced by randomly choosing 7725 genes from non-target genes. The data frame for the predictors was previously described (Inagaki et al, 2021) (https://github.com/soinagak/FLD2021). The RNAPII Ser2P ratio and Ser5P ratio were added to predictor variables of this data frame.

The random forest was trained with genes on chromosomes 1–4. Cross-validation was performed with genes on chromosome 5. R package randomForest (https://www.rdocumentation.org/packages/randomForest/versions/4.6-14/topics/randomForest) was used with option ntree = 1,000. The ROC curve was plotted using R package ROCR (Sing et al., 2005).

### Co-immunoprecipitation (Co-IP)

The Co-IP experiments were performed as described in (Pfab et al, 2017) with minor modifications. 2 grams of 14-day-old seedlings were used for a Co-IP. The sample was ground into fine powder and suspended in 10 ml of Extraction buffer (25 mM HEPES-KOH pH 7.4, 100 mM NaCl, 2 mM MgCl2, 10 % glycerol, 0.05 % IGEPAL CA-630, 1 mM DTT, 5 mM EGTA, 1x cOmplete proteinase inhibitor). MgCl2 was added to a final concentration of 5 mM and 50 U/mL benzonase was added to the mixture. The mixture was incubated for 30 min at 4 ºC on a rotating wheel and pelleted by centrifugation at 4,000 g for 10 min at 4ºC. The supernatant was filtered through a 40 μm nylon cell strainer on ice. IP was performed from the supernatant using FLAG antibody-conjugated beads or only beads (mock) for 1 h. The beads were washed with 1 mL of ice-cold Extraction buffer three times.

### Western blotting

Bulk histones for western blotting were prepared using Histone extraction kit (Active Motif). RNAPII samples for western blotting were prepared as described in the method for chrRNA-seq. 50 μl of chromatin suspension were mixed with an equal volume of 2X SDS buffer and boiled for 5 minutes at 95 ºC.

Proteins were resolved on SDS-PAGE, and transferred to a PVDF membrane using the Trans-Blot Turbo and Trans-Blot transfer pack (Bio-Rad). The primary and secondary antibody reactions were performed using an iBind Flex Western System (Thermo Fisher Scientific). Primary antibody (H3K4me2 (ab32356; Abcam), H3 (ab1791; Abcam), RNAPII total CTD (Diagenode, C15100055), RNAPII phospho S2 (MABI0602; MBL), RNAPII phospho S5 (MABI0603; MBL), FLAG (F1804; SIGMA) and secondary antibody (Anti-Rabbit IgG HRP (NA934; Cytiva), Anti-mouse IgG HRP (NA931, Cytiva)) reactions were performed at room temperature for more than 2.5h according to the manufacturer’s protocol. The blots were developed with ECL prime solution (Cytiva), and the signal was quantified by iBright Imager (Thermo Fisher Scientific).

### RT-qPCR

Total RNA was isolated from aerial parts of three 14-day-old seedlings grown on MS media, using the RNeasy Plant Mini Kit (Qiagen) and cDNA was synthesized using ReverTra Ace qPCR RT Master Mix with gDNA Remover (TOYOBO). Real-time PCR was performed using LightCycler 96 (Roche) and THUNDERBIRD SYBR qPCR mix (Toyobo) with 50 cycles of denaturation at 95 °C for 15 s and extension at 60 °C for 1 min using specific primers for LDL3 (forward: TCCTAGTTGGAAGCATGACCAACGAGCG reverse: GAGTCGAGGAACTGTGTAACAGCCATAGC), and for 60S ribosomal protein gene AT3G49010 (forward: TCGCAAGAACCGATCTTTGGAGG, reverse: AACTCTTCTGGTGTAGAGTCACC) as a reference. The plant tissues for gene expression analysis were produced in three biological replicates and three technical replicates of each repetition were carried out.

## Data Availability

The high-throughput sequencing data generated in this study is available in the NCBI database under the accession number PRJNA934737.

## Acknowledgements

We thank all Kakutani laboratory members and especially Dr. Taku Sasaki and Dr. Akihisa Osakabe for helpful discussion, advice, and technical assistance. This work used the Vincent J. Coates Genomics Sequencing Laboratory at UC Berkeley, supported by NIH S10 OD018174 Instrumentation Grant. The computations were partially performed on the NIG supercomputer at NIG, Japan. We thank NASC/ ABRC for distributing the seeds. This work was supported by grants from Japan Science and Technology Agency (JST) PRESTO (no. JPMJPR17Q1) to S.I., JST CREST (no. JPMJCR15O1) and HFSP (RGP0025/2021) to T.K., and Japan Society for the Promotion of Science (JSPS) (nos. JP26221105, JP15H05963 and JP19H00995 to T.K., JP20H05913 and JP22H02299 to S.I.).

## Author Contributions

S.M., S.O., S.I., and T.K. designed the study. S.M., S.O., M.T., and K.T. performed the experiments. S.M. analyzed the data. S.M., S.I., and T.K. wrote the paper with incorporating comments from the other authors.

## Figure legends

**Appendix Figure S1. LDL3 has the same demethylation target genes across several tissues. (**A, B) H3K4me2 levels in roots (A) and callus (B) of the *ldl3* mutants compared with WT (data from Ishihara et al, 2019). Each dot represents RPKM within each protein-coding gene. Red dots, protein-coding genes with hyper H3K4me2 in shoot tissues of *ldl3* (7,725 genes; Fig. 1A).

**Appendix Figure S2. H2Bub and H3K36me3 do not affect H3K4me2 demethylation by LDL3**. (A) Western blotting of H2B and H2Bub on bulk histone extracted from the *hub* mutants. (B) Averaged profiles of H3K4me2 around LDL3 target genes in each genotype (WT, *hub1, hub2, sdg8*). The numbers of genes analyzed are 7,367 as in Fig. 2A. The ribbons indicate s.e.m. (C) H3K4me2 levels in each of the mutants compared with WT. Each dot represents RPKM within each protein-coding gene. Red dots, protein-coding genes with hyper H3K4me2 in *ldl3* (2,809 genes; Fig. 2C).

**Appendix Figure S3. Defects in RNAPII phosphorylation and transcriptional elongation mimic loss of LDL3 function**. (A) Averaged profiles of H3K4me2 changes around LDL3 target genes in the mutants compared to WT. The numbers of genes analyzed are 7,367 as in Fig. 2A. The ribbons indicate s.e.m. (B) A biological replicate of the experiment shown in Fig. 2B. The protein-coding genes were sorted in the same way as Fig. 2B.

**Appendix Figure S4. Paf1C component mutant plants showed increases of H3K4me2 in genes demethylated by LDL3**. H3K4me2 levels in each mutant compared with WT. Each dot represents the RPM within each protein-coding gene. Red dots, protein-coding genes with hyper H3K4me2 in *ldl3* (H3K4me2 levels in *ldl3* (RPM) - H3K4me2 levels in *ldl3* (RPM) > 5; 8,967 genes). The effects of mutations in Paf1C components on the phenotype and H3K4me2 levels appeared to be correlated (Tamada et al, 2009; Yu & Michaels, 2010).

**Appendix Figure S5. H2Bub and H3K26me2/me3 are controlled downstream of Paf1C**. (A-C) Averaged profiles of H2Bub (A), H3K36me2 (Bb) and H3K36me3 (C) around LDL3 target genes (n=7367; Fig. 2A) and the other protein-coding genes (n=19,839) in each genotype (WT, *ldl3, elf8*). The ribbons indicate s.e.m.

**Appendix Figure S6. H3K4me1/me3 in *ldl3, elf8*, and *cdkf;1* mutants**. (A) H3K4me1/me3 levels in each of the mutants compared with WT. Each dot represents the RPKM within each protein-coding gene. Red dots, protein-coding genes with hyper H3K4me2 in *ldl3* (7,367 genes; Fig. 2A). (B) Averaged profiles of H3K4me1/me3 around LDL3 target genes (n = 7367; Fig. 2A) in each genotype (WT, *ldl3, elf8, cdkf;1*). The ribbons indicate s.e.m. (C) Relationships between changes in H3, H3K4me1, and H3K4me3 levels (RPKM) in *elf8* and *cdkf;1*, and in *ldl3* within each protein-coding gene compared to WT.

**Appendix Figure S7. Paf1C and CDKF;1 do not largely affect the transcript level of LDL3**. mRNA level of *LDL3* gene in WT, *ldl3, cdkf;1*, and *elf8*. Relative transcript level determined with RT-qPCR was normalized to the internal control (60 S ribosomal protein coding gene). Means and SD for three biological replicates are shown.

**Appendix Figure S8. The *elf8* and *cdkf;1* mutants compromise shoot regeneration from callus similar to the *ldl3* mutant**. (A, B) Shoot regeneration phenotype in each genotype (WT, *ldl3, elf8, cdkf;1*). The shoot regeneration rate was reduced in *ldl3, elf8, cdkf;1*. Root tip explants were incubated on CIM 14 days (A), and on SIM 14 days (B). Scale bar: 10 mm

**Appendix Figure S9. RNAPII dynamics was not largely affected in the *ldl3* mutant** (A) Averaged profiles of RNAPII (Ser5P/total CTD ratio) in the gene body around LDL3 target genes (n = 7367; Fig. 2A) for WT, *ldl3, elf8*, and *cdkf;1*. The ribbons indicate s.e.m. (B) Heatmaps of RNAPII Ser5P levels (Ser5P/total CTD ratio) are shown in WT and the three mutants. The protein-coding genes were sorted based on increases in gene-body H3K4me2 levels in *ldl3*. (C) Averaged profiles of RNAPII (Ser5P/total CTD ratio) in the gene body around LDL3 target genes for WT and *ldl3*. The ribbons indicate s.e.m. (D) Averaged profiles of chrRNA around LDL3 target genes (n=7367; left) and all protein-coding genes (n=27,206; right) for WT and *ldl3*. The ribbons indicate s.e.m.

**Appendix Figure S10. The accumulation of H3K4me2 in the *elf8* mutant is mostly mediated by *LDL3***. (A) Box plots of H3K4me2 levels (RPM) in WT and each of the mutants in eight different subsets of genes, grouped by k-means clustering. Genes were classified into eight clusters based on H3K4me2 levels in wild-type, *elf8-1, elf8-2, ldl3* and *ldl3 elf8*. Cluster 4 (12,731 genes) is a group of genes whose H3K4me2 levels are unchanged in mutants, whereas genes in clusters 1, 3 and 5 have elevated H3K4me2 levels in *ldl3* and *elf8* mutants. The *ldl3 elf8* double mutant had an additional increase of H3K4me2 levels than that in the *ldl3* single mutant in some genes (797 genes in cluster 5), but in the majority of genes (5,146 genes in clusters 1 and 3), the change in H3K4me2 levels in *ldl3 elf8* double mutant were similar to that of the *ldl3* mutant, supporting the idea that *ELF8* and *LDL3* work mainly in the same pathway. (B) The *ldl3* mutant has wild type appearance, but the *ldl3* mutation enhances the *elf8* mutant phenotype.

**Appendix Figure S11. Correlations between H3K4me and transcription of protein-coding genes in WT, *elf8*, and *cdkf;1***. Protein-coding genes are ranked by H3K4me1, me2 or me3 (x-axis, RPKM) and expression levels (y-axis, chrRNA-seq RPKM). The data of WT is the same as Fig. 6A. The densities of genes are visualized as heat maps. ρ: Spearman’s correlation coefficient.

**Appendix Figure S12. Amino acid alignment of the SRI domain in LDL3 and its homologs**. Regular secondary structure of the SRI domains in H. sapiens SETD2 is reported above the alignment. The SRI domain forms a closed three-helix bundle (α1, α2, α3). In the alignment, positions conserved or highly similar among the entire SRI family are colored in green for hydrophobic amino acids (VILM), orange for aromatic amino acids (FYW), blue for basic (KRH) amino acids and pink for acidic (DE) amino acids. Host species, NCBI accession number and domain limits (in brackets) are indicated at the start of the sequences.

